# Glioblastoma utilizes fatty acids and ketone bodies for growth allowing progression during ketogenic diet therapy

**DOI:** 10.1101/659474

**Authors:** Jantzen Sperry, Michael C. Condro, Lea Guo, Daniel Braas, Nathan Vanderveer-Harris, Kristen K.O. Kim, Whitney B. Pope, Ajit S. Divakaruni, Albert Lai, Heather Christofk, Maria G. Castro, Pedro R. Lowenstein, Janel E. Le Belle, Harley I. Kornblum

## Abstract

Glioblastoma (GBM) metabolism has traditionally been characterized by a primary dependence on aerobic glycolysis, prompting the use of the ketogenic diet (KD) as a potential therapy. In this study we evaluated the effectiveness of the KD in GBM and assessed the role of fatty acid oxidation (FAO) in promoting GBM propagation. *In vitro* assays revealed FA utilization throughout the GBM metabolome, and growth inhibition in nearly every cell line in a broad spectrum of patient-derived glioma cells treated with FAO inhibitors. *In vivo* assessments revealed that knockdown of carnitine palmitoyltransferase 1A (CPT1A), the rate limiting enzyme for FAO, reduced the rate of tumor growth and increased survival. However, the unrestricted ketogenic diet did not reduce tumor growth, and for some models significantly reduced survival. Altogether, these data highlight important roles for FA and ketone body metabolism that could serve to improve targeted therapies in GBM.

## Introduction

Normally, the adult brain meets most of its energy demand by complete oxidation of glucose, producing pyruvate which is converted into acetyl-CoA for entrance into the TCA cycle to support the electron transport chain (Agnihotri and Zadeh, 2016). Thus, glycolysis and respiration remain tightly linked, resulting in efficient ATP production with little lactic acid production. On the other hand, many tumors, including the universally lethal glioblastoma (GBM), utilize the Warburg effect (Vander Heiden et al., 2009) which dramatically increases the rate of aerobic glycolysis and lactate production in the cytosol. This process allows for increased biomass production and facilitated invasion through acidification of the microenvironment amongst other things.

The signaling pathways hyperactivated in GBM and the underlying mutations significantly influence GBM metabolism. Numerous oncogenic mutations activate the PI3K/AKT/mTOR signaling pathway which promotes glucose utilization and the Warburg effect (Yang et al., 2019). The net result of these mutations is to both utilize glucose as an energy substrate and to promote FA synthesis for membrane biogenesis (Furuta et al., 2008). Thus, GBM-causing mutations induce what has been called a “glucose addiction”, which one might target directly by limiting glucose availability. The effects of some other mutations, such as those in the isocitrate dehydrogenase (IDH) 1 and 2 genes on glucose utilization and dependence are less clear.

Brain neurons utilize glucose as a primary energy source under normal physiological conditions, and upon glucose deprivation neurons readily utilize ketone bodies (KBs) as an alternative fuel (Maurer et al., 2011). Glial cells are also capable of using KBs and FAs for fuel (Lopes-Cardozo et al., 1986; Weightman Potter et al., 2019). Thus, one might reason that treatments that inhibit glucose metabolism and promote FA and KB metabolism would be ideal for treating tumors that are known to be glucose addicted. Indeed, the ketogenic diet (KD), which consists of high fat, low carbohydrates and adequate protein, has long been used in the treatment of epilepsy, and has recently been proposed as a potential adjuvant therapy for a number of cancers including GBM (Maurer et al., 2011; Vidali et al., 2015). In some animal studies, the KD has indeed shown preliminary efficacy (Abdelwahab et al., 2012; Mukherjee et al., 2019) and a number of glioma patients have initiated KD therapy.

Despite the potential for utilizing the KD as a therapy for glioma, there are factors that may mitigate enthusiasm. First, not all animal studies using the KD have demonstrated efficacy (De Feyter et al., 2016). Another factor to consider is glioma heterogeneity. In addition to there being multiple molecular subtypes of GBM (Lee et al., 2018), with some mutations that may not create glucose addiction, there is a high degree of intratumoral heterogeneity. In some regions of GBM there is relative hypoxia which could alter the tumor’s ability to utilize glucose via the Warburg effect (Kim et al., 2006). These properties may be exacerbated by therapies that limit vascular supply, selecting for those cells most capable of surviving under harsh conditions. Recent studies have indicated that GBMs could be utilizing FAs as both energy sources and as substrates for generating cellular building blocks (Strickland and Stoll, 2017). However, it is unknown whether FAO occurs across the mutational spectrum of GBM. Nor has it been demonstrated precisely how FAs are utilized, or whether the KD itself may supply FAs and/or KBs.

Here we demonstrate that the well-characterized U87 glioma cell line and patient-derived GBM cultures, including those with mutations in the PI3K/AKT/mTOR pathways and the IDH1 gene, utilize FAs and KBs for growth under physiologic and standard cell culture conditions. Metabolic tracing studies indicate FAs are oxidized and incorporated into TCA cycle intermediates. Furthermore, administration of the non-calorie restricted KD to tumor-bearing animals does not decrease the rate of tumor growth, nor does it improve animal survival. Knockdown of CPT1A, the rate-limiting enzyme for FAO, or its inhibition by etomoxir alters the metabolic profile and growth of GBM *in vitro*. These data highlight the metabolic plasticity of GBM and caution against universal use of the KD as adjuvant therapy, although they do not preclude combining the KD with other metabolic therapies and do not address the potential for calorie-restricted KD utilization.

## Results

### Fatty acid and ketone oxidation in glioma cells

In order to determine whether GBMs are capable of oxidizing fatty acids we first examined expression levels of *CPT1,* the rate-limiting enzyme required for FAO, in U87 cells. While the precise origin of this cell line is not known, a recent study has validated it as a bona fide glioblastoma line (Allen et al., 2016). and it is widely used as a model in glioma research, especially in metabolic studies including studies of fatty acid metabolism (Guo et al., 2009). Analysis of the TCGA dataset from GBM tumors reveals that of the three *CPT1* isoforms found in humans, *CPT1A* and *CPT1C* are most highly expressed in GBM, and that *CPT1B* is expressed at lower levels (Figure 1A). Because the TCGA data are derived from whole tumors, we utilized a single cell RNA seq dataset (Darmanis et al., 2017) to verify the expression of CPT isoforms, as well as the major enzymes involved in fatty acid beta oxidation, within tumor cells themselves, and that the levels of expression of these genes were comparable to those of fatty acid synthesis. (Figure S1).

**Figure 1.**
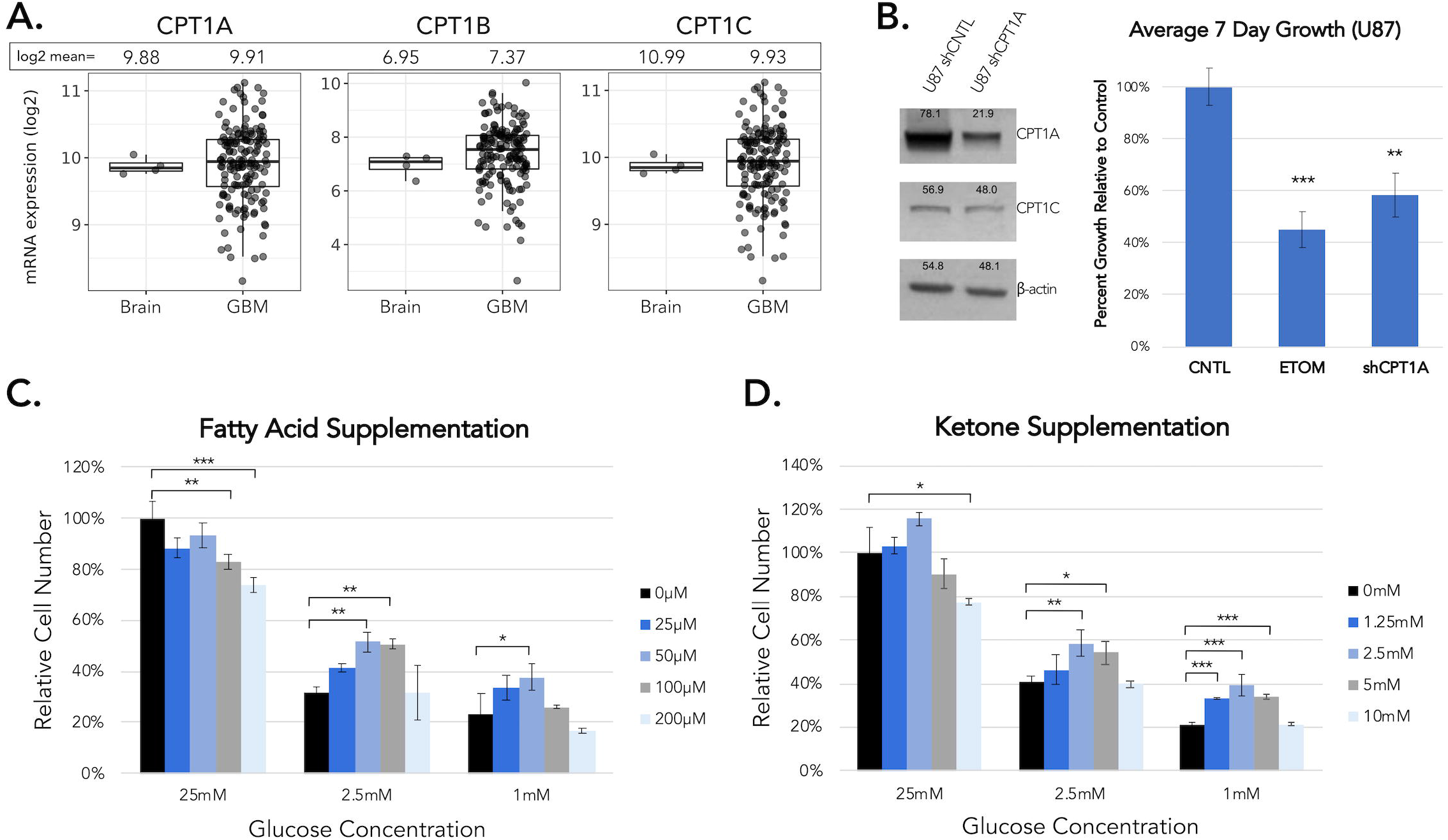
U87 GBM cells oxidize fatty acids and ketone bodies to support growth. **A.** Relative mRNA expression of the three CPT1 isoforms in GBM based on analysis of the TCGA dataset from patient-derived GBM tumors. N = 4 non-tumor, N = 156 GBM. See also Figure S1. **B.** Quantification of the effects of 100 μM ETOM treatment and shRNA targeting CPT1A on U87 cell growth. Culture growth was assessed following a 7 day treatment with ETOM administered twice per week. n = 3 replicates. Error bars indicate ± SD. Univariate generalized linear model F(2,5) = 47.37, p = 5.6×10^−4^, post-hoc Tukey HSD (** = p<0.01; *** = p<0.001). **C**. Effects of FA and ketone supplementation under variable concentrations of glucose following a 7 day growth period in U87 cells. n = 3 replicates. For all univariate generalized linear models for different concentrations of supplements within each glucose concentration p<0.01. Post-hoc Tukey HSD significance is reported for each supplement concentration against its control. Error bars denote ± SD. (* = p<0.05; ** = p<0.01; *** = p<0.001).

To assess a potential role for FAO in gliomas, we treated U87 GBM cells with the CPT1-specific inhibitor etomoxir (ETOM) *in vitro* and found that it modestly inhibited growth under standard, serum-free conditions (Figure 1B). Additionally, shRNA-mediated partial (68%) knockdown of CPT1A alone, was sufficient to significantly inhibit *in vitro* cell growth by ~40% to levels near those observed with ETOM alone (Figure 1B), suggesting that the CPT1A isoform predominantly contributes to FAO in these cells. Importantly, this demonstrates that endogenous FAO contributes to the maintenance of normal *in vitro* cell growth in these GBM cells despite U87s having a PTEN mutation with increased PI3K/AKT/mTOR signaling, which has been previously thought to inhibit FAO in GBM. Because gliomasphere media used for *in vitro* studies contains levels of glucose (25 mM) not likely to be found naturally in the human brain or in brain tumors, a glucose-limiting assay was performed to assess both fatty acid and ketone body utilization under more physiologically relevant glucose levels, with 2.5 mM glucose representing extracellular levels commonly found in the brain and 1 mM glucose serving as a low glucose condition to more closely mimic levels found in tissue that do not have adequate access to vascular supplies of glucose (Silver and Erecińska, 1994). Palmitic acid supplementation (50-100 μM) increased U87 cell growth under 2.5 mM low glucose conditions (Figure 1C), whereas ketone supplementation with 2.5-5 mM 3-hydroxybutyrate (3-OHB) promoted cell growth in both 2.5 mM and 1 mM glucose (Figure 1D). Higher levels of supplementation (200 μM palmitic acid and 10mM 3-OHB) resulted in a reversal of the proliferative effects, likely due to toxicity of these supplements at higher concentrations, as has been described for neural stem cells and another tumor cell line (Park et al., 2011; Poff et al., 2014). Neither FA nor ketone supplementation had growth-promoting effects under non-physiological conditions of high glucose (25 mM).

To further determine how and where FAs are utilized within U87 cells, untreated CNTL cells, cells treated with ETOM, and cells with CPT1A knockdown were subjected to LC-MS metabolomic analysis using fully-labeled ^13^C palmitic acid in physiological glucose conditions (2.5 mM). Analyzing fractional contribution (i.e. the percent contribution of carbons in each metabolite from radiolabeled palmitate) within these samples revealed labeling throughout the TCA cycle as well as in the ubiquitously important currency metabolite ATP, indicating that FAO contributed to the pool of carbons used in the synthesis of these metabolites. Treatment with ETOM or CRISPR-based CPT1A knockdown (Figure S2), dramatically diminished this labeling (Figure 2A). Similar effects were observed when analyzing the relative amounts of these metabolites, wherein ETOM treatment and CPT1A knockdown significantly decrease levels of TCA-associated metabolites, acetyl-CoA, and ATP (Figure 2B). The effects of FAO inhibition on relative amounts of metabolites demonstrates that inhibiting FAO affects production of these metabolites metabolites via unclear mechanisms, as the oxidation of even chain fatty acids should not make a net contribution to carbons in intermediary metabolites. Taken together these data confirm that GBM cells actively oxidize FAs and suggest that of the three CPT1 isoforms, CPT1A appears to be the primary isoform regulating this process in the U87 line.

**Figure 2.**
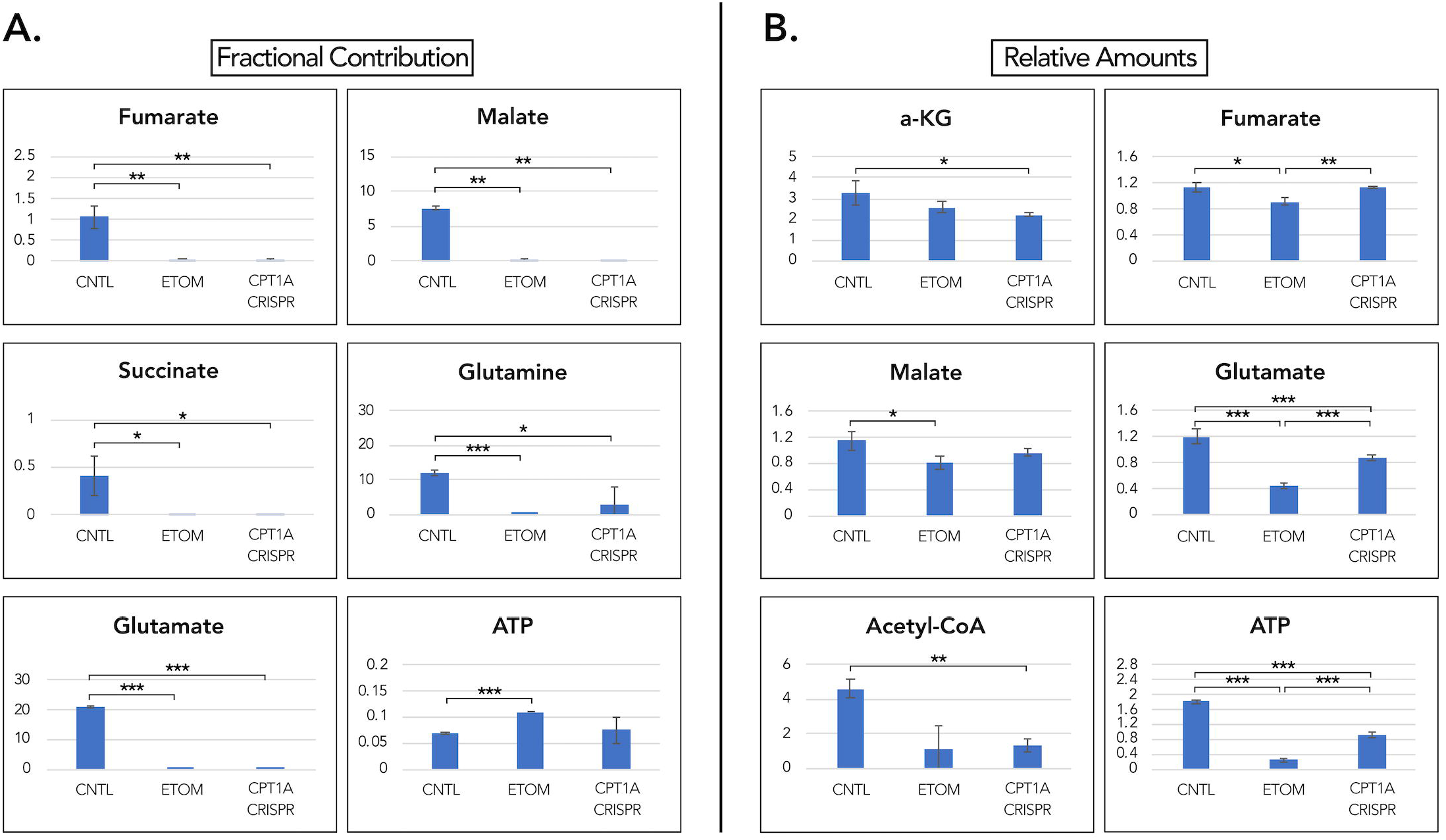
U87 GBM cells oxidize fatty acids to generate TCA cycle intermediates and currency metabolites. **A-B.** Effects of ETOM and CPT1A knockdown in U87 cells on fractional contribution (the percent contribution of carbons from radiolabeled palmitate) and relative amounts of relevant TCA cycle intermediates and currency metabolites using ^13^C-palmitate LC-MS analysis. Glucose concentration = 2.5 mM. n = 3 replicates. Error bars = ± SD. (*=p<0.05, **=p<0.01, ***=p<0.001). See also Figure S2.

### CPT1A knockdown and the ketogenic diet

Based on these initial results and previous studies (Seyfried et al., 2003; Skinner et al., 2009) that have indicated that many brain tumors may lack the ability to oxidize ketones as an energy substrate, we sought to determine if inhibiting FAO while concurrently implementing a ketogenic diet using an orthotopic xenograft model in mice might serve as an effective therapeutic strategy. It is thought that this strategy would reduce the availability of glucose as an energy substrate and make the GBM cells more vulnerable to cell death (starvation) by also inhibiting FAO as an alternative energy source. To assess these results *in vivo,* as well as to determine the effectiveness of the KD on tumor growth, U87 cells expressing a firefly luciferase-GFP reporter (FLuc-GFP) and either shCNTL or shCPT1A were implanted unilaterally into the striatum of adult NSG mice. Four days after implantation, half of each group was switched to a calorie-unrestricted KD followed by weekly bioluminescent imaging with luciferin (Figure 3A). At 21 days post-injection, animal weights, blood glucose levels, and blood ketone levels were evaluated. Both of the ketogenic diet cohorts (shCNTL and shCPT1A) exhibited decreased total body weight, decreased blood glucose levels, and increased blood 3-OHB levels, confirming ketosis was achieved in these animals (Figure 3B). Contrary to our initial hypothesis, analysis from weekly bioluminescent imaging shows that the shCNTL cohort receiving the KD had the greatest increase in tumor size, although the difference between this group and the shCNTL standard diet (STD) group was not significant (Figure 3C). Survival analysis also indicated that this group had the poorest survival (Figure 3D), supporting the hypothesis that the KD contributed to tumor growth in this cohort. Importantly the growth of the tumors treated with the KD was significantly diminished (Figure 3C) and survival enhanced (Figure 3D) by CPT1A knockdown. CPT1A knockdown resulted in a decrease in tumor growth and enhanced survival in the mice on the standard diet as well, suggesting that FAO contributes to tumor growth even under normal dietary conditions. Immunohistochemisty of sectioned post-mortem tissue confirmed that CPT1A knockdown was maintained throughout the length of the experiment (Figure 3E). The findings using our model, while surprising, may indicate that at the very least ketosis does not hinder GBM growth, and may exacerbate it in some tumors, while also highlighting an important role of CPT1A-mediated FAO in tumor growth in general.

**Figure 3.**
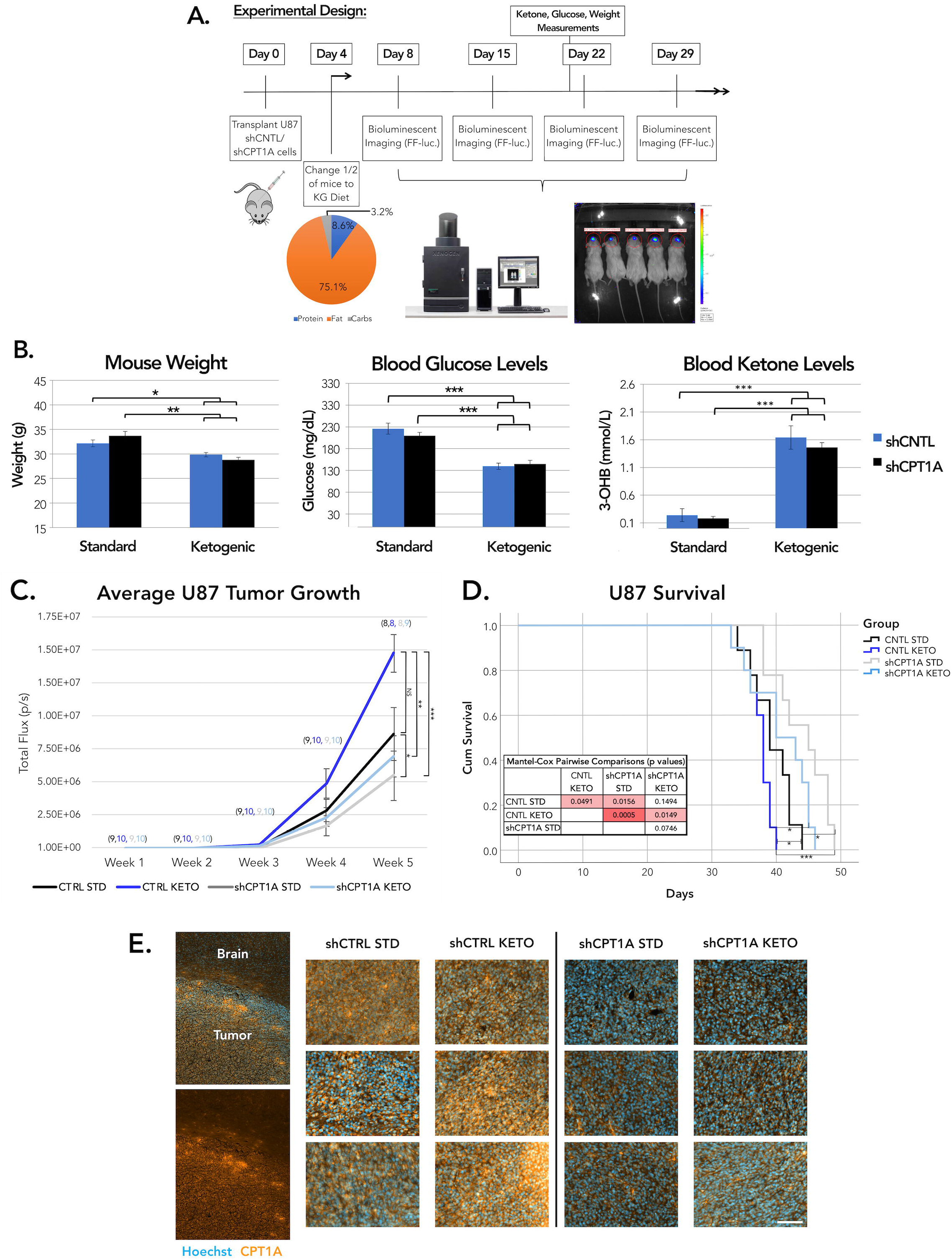
Knockdown of CPT1A, but not ketogenic diet, decreases U87 tumor growth *in vivo.* **A.** Basic schematic of the experimental design for the U87 shCPT1A ketogenic diet xenotransplant experiment. Cells were implanted into 10 mice per group on Day 0. Mice were kept on standard diet for 4 days to allow appropriate recovery, after which half of each group was placed on the ketogenic diet. Weekly bioluminescent imaging was initiated on Day 8. **B.** On Day 21 mouse weight, blood glucose and ketones were measured. Error bars denote ± SEM. Mixed effects model F(*3,55.111*) = 4.401, p= 0.0076. Last time point F(*3,29*) = 5.618, p = 0.0037, pairwise comparison signifance values are noted in the figure. **C.** Average tumor growth based on total flux (photons/s) from the firefly luceriferase. (Error Bars = ± SEM). Numbers in parentheses indicate the number of animals measured at each time point. Mixed effects model with repeated measures F(*1,4,3*) = 4.401, p = 0.008, followed by pairwise post-hoc Tukey HSD comparisons at the final time point. (* = p<0.05, ** = p<0.01, *** = p<0.001). **D.** Kaplan-Meier curve of mouse survival over the course of the experiment. Survival was tracked for individual animals until either death or progression to a moribund state. **E.** Representative immunohistochemistry of CPT1A in sectioned tissue following brain perfusion. Each image represents a different animal, 3 animals per group. Scale bar = 100μm.

### Patient-derived GBM gliomasphere cultures

To further assess inhibition of endogenous FAO on GBM cell growth and survival we tested the effects of ETOM in a panel of patient-derived GBM cultures. These experiments were done under high glucose conditions with the minimal exogenous fatty acids present in B27 supplement. A seven day treatment with ETOM resulted in diminished overall culture growth for all gliomasphere lines tested (Figure 4A). This analysis yielded no significant differences with regard to TCGA classification, IDH1 mutant status, PTEN deficiency, EGFR amplification, or the presence of the EGFRviii mutation (data not shown). Interestingly, ETOM treatment did not inhibit the growth of cultured human fetal neurospheres or SVZ-derived mouse neurospheres. This may indicate that FAO is GBM-specific, irrespective of mutational status, and less important to healthy tissue, making it a potentially attractive therapeutic target. A seven day growth assay in a subset of patient-derived GBM lines of varying mutational status cultured a physiologically relevant level of gluose (2.5 mM) showed a similar sensitivity to ETOM inhibition of endogenous FAO to what was observed in 25 mM glucose (Figure S3). The effects of ETOM were comparable to CPT1A knockdown in a highly sensitive line, HK301 (Figure 4B). Following a four day treatment with ETOM we found a significant decrease in actively dividing cells as assessed using bromodeoxyuridine (BrdU) incorporation (Figure 4C), demonstrating the importance of endogenous FAO to the maintenance of cell proliferation even in a GBM line with a mutation that enhances AKT pathway signaling (HK301). In addition to these effects on proliferation we also observed an increase in cell death as measured by annexin V/PI flow cytometry following treatment with ETOM (Figure 4D). Additionally we examined the effect of FA and 3-OHB supplementation on growth of HK157 in 25 mM, 2.5 mM, and 1 mM glucose (Figure 4E,F). Similar to U87 cells (Figure 1C-D), a significant increase in growth was observed with FA and 3-OHB supplementation in physiologically relevant (2.5 mM) glucose. Taken together, these data support our hypothesis that FAO is a significant contributor to overall GBM metabolism and growth, which is not prevented by PI3K/AKT/mTOR pathway-promoting mutations in our lines, as has been observed in previous studies on glucose dependency and fatty acid utilization in GBM (Buzzai et al., 2005; Yang et al., 2009; Ye et al., 2013).

**Figure 4.**
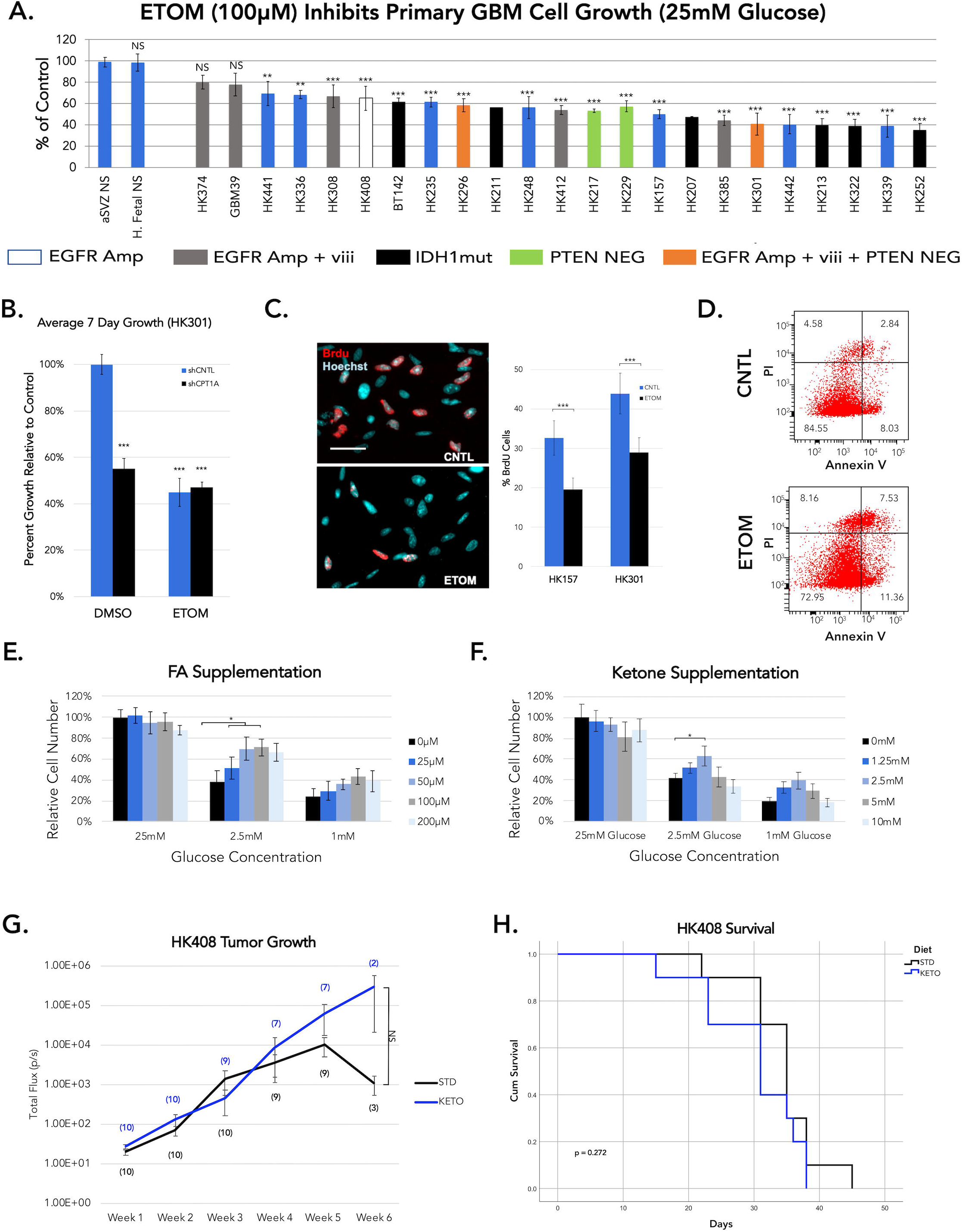
Etomoxir inhibits patient-derived GBM cell proliferation and promotes cell death. **A.** Quantification of the effects of 7 day 100 μM ETOM treatment administered twice per week across a panel of patient-derived GBM cell lines and two non-GBM controls (adult mouse-derived neurospheres and human fetal-derived neurospheres) relative to vehicle-treated controls. n = 3 replicates, except for HK211 and HK207 in which only a single measurement was taken. Error bars = ± SEM. Significance values are relative to aSVZ NS with Bonferroni correction for multiple comparisons. Cultures that are EGFR amplified, EGFR ampified + EGFRviii mutant, IDH1 mutant, and PTEN deficient are indicated. See also Figure S3. **B**. Average 7 day growth of patient-derived cell line HK301with both CPT1A knockdown and 100 μM ETOM. n = 3 replicates. Univariate generalized linear model F(3,8) = 99.8, p = 1.12×10^−6^, post-hoc Tukey HSD relative to control (*** = p<0.001). **C.** Assessment of actively dividing cells by BrdU immunocytochemisty in two patient-derived GBM lines grown for 4 days in the presence of 100 μM ETOM. n = 9 replicates. Error bars = ± SEM. **D.** Flow cytometry analysis of cell death using dual Annexin V/PI staining in HK157 cells following 4 day treatment with 100 μM ETOM. **E.** Effects of pamitic acid or **F.** 3-OHB supplementation at different glucose concentrations on growth of HK157 cells *in vitro.* n = 3 replicates. Error bars = SD. Significance values represent T-tests of each condition compared to the control for the same concentration of glucose (* = p<0.05). **G.** Tumor growth measured by bioluminescence imaging for patient-derived cell line HK408. The effect of diet was not statistically significant by mixed effects model (p>0.05). Numbers adjacent to lines indicate N at each time point. **H.** Kaplan-Meier curves for the data in E. Diet had no significant effect on survival by Mantel-Cox log-rank values (p = 0.272). See also Figure S4.

To determine the effects of our non-calorie-restricted KD on a primary patient-derived cell xenograft model *in vivo*, we implanted a modestly ETOM-sensitive line, HK408 into the striatum and placed the mice on the KD or standard diet after two days. There were no significant differences in tumor growth or animal survival between the two diets. (Figure 4G, 4H). To assess the effects of the KD in an immune competent model, we utilized a syngeneic murine model (Núñez et al., 2019). Again, we found no significant benefit of the KD in this model (Figure S4). Thus, of the three cell lines utilized we did not find a therapeutic benefit of the KD in any of the models tested.

### LC-MS with fully-labeled palmitic acid in a primary cell line

We utilized LC-MS analysis with fully-labeled ^13^C palmitic acid to characterize the contribution of FAO to the metabolome of patient-derived GBM cells of a different, moderately ETOM sensitive cell line, HK157. Fractional contribution studies revealed that ETOM treatment caused widespread changes in carbon labeling across the metabolome (Figure 5A). Importantly, ETOM resulted in in diminished labeling of acetyl-CoA, the end product of FAO, along with an increase in the amount of labeled palmitate within the cell (Figure 5B). Both results confirmed that labeled FAs were taken up and oxidized within the cell. In addition to acetyl-CoA, ETOM treatment also resulted in diminished labeling within the TCA cycle and among metabolites that commonly feed into the TCA cycle such as glutamine, glutamate, and lactate (Figure 5B). Analysis of the relative amounts of metabolites with ETOM treatment did not result in changes to overall acetyl-CoA or palmitate levels. However, ETOM treatment did result in lower relative levels of TCA-associated metabolites which include alpha-ketoglutarate, citrate, fumarate, glutamine, and glutamate (Figure 5C). A full list of significantly altered metabolites resulting from FAO inhibition with ETOM, with fractional contribution shown in Table 1 and relative amounts shown in Table 2, demonstrates that GBM cells utilize FAO to support a number of critical metabolic pathways throughout the GBM metabolome.

**Figure 5.**
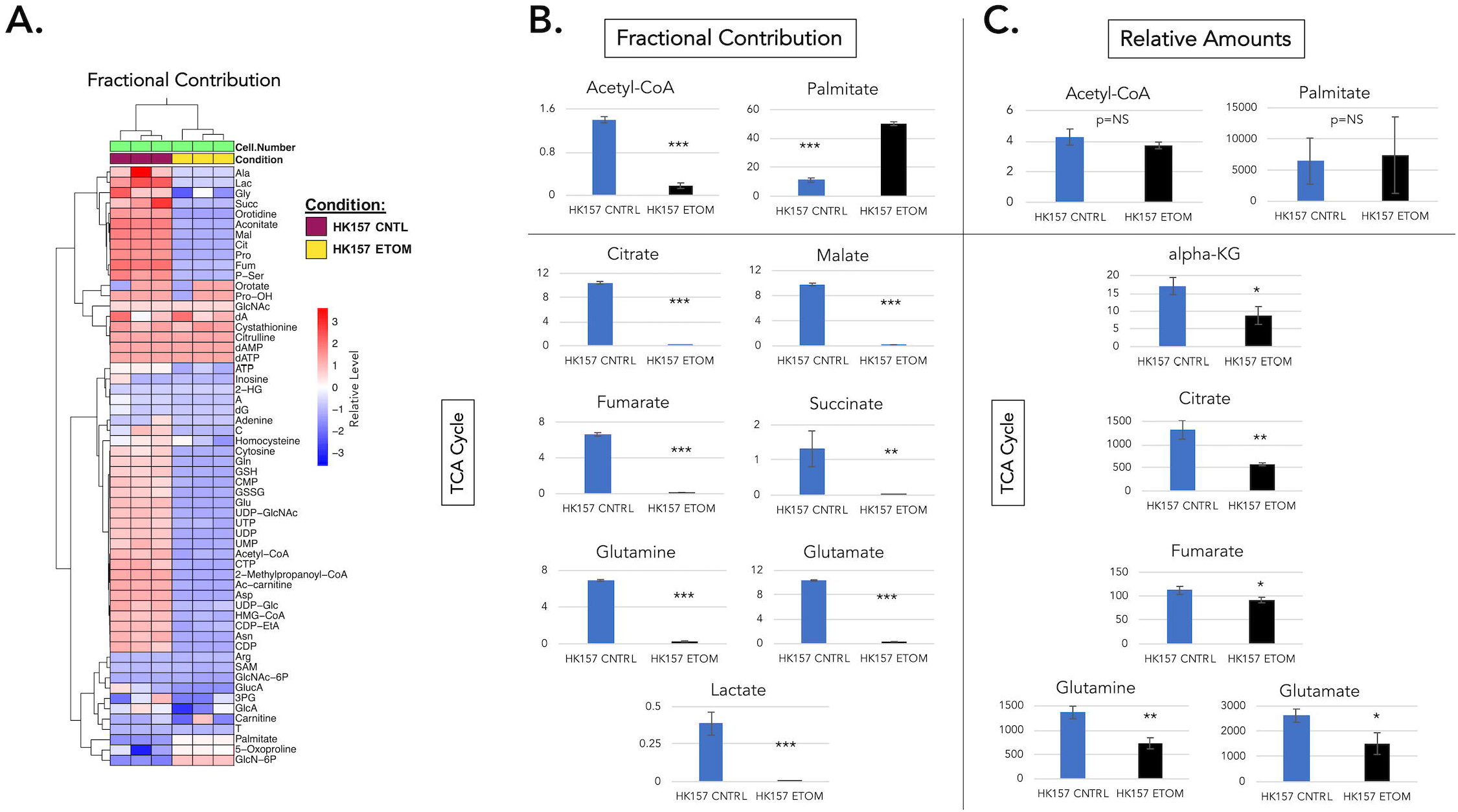
GBM cells oxidize FAs to generate acetyl-CoA for use within the TCA cycle. **A-B.** LC-MS analysis of ^13^C-palmitic acid fractional contribution following treatment with 100 μM ETOM. **A.** Heat map showing effects of ETOM on fractional contribution across metabolites. Relative expression is measured as standard deviations from the group mean. **B.** Quantification of the effects of ETOM on intracellular carbon labeling of acetyl-CoA, palmitic acid, and other relevant metabolites associated with the TCA cycle. **C.** Effects of ETOM on relative amounts of relevant metabolites as in B. ETOM concentration used: 100 μM. n = 3 replicates. Error bars = ± SD. (* = p<0.05, ** = p<0.01, *** = p<0.001).

**Table 1.**
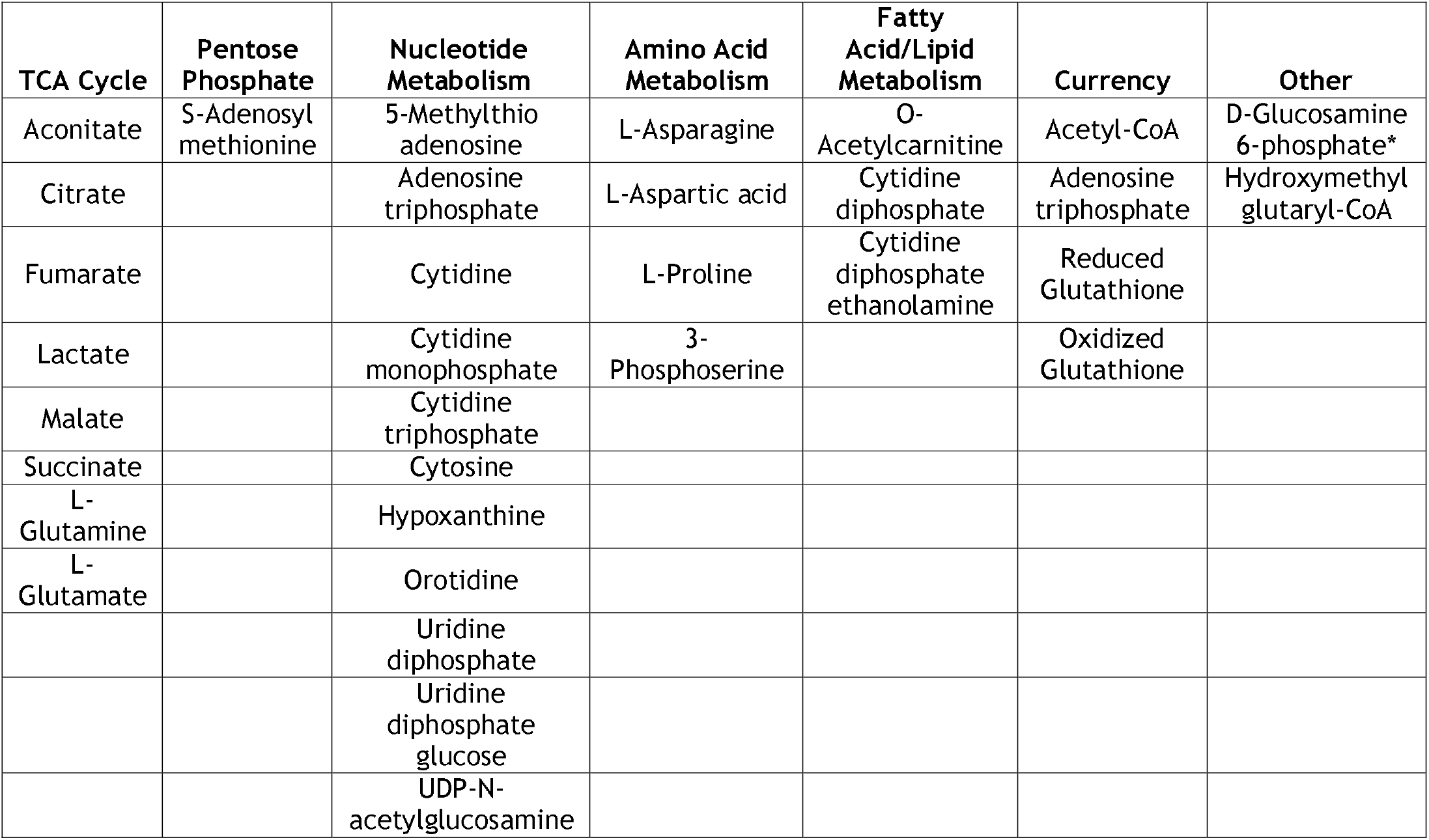
List of all metabolites altered by ETOM (100 μM) treatment with regard to fractional contribution for HK157. Metabolites were measured by labeled ^13^C-palmitic acid using LC-MS analysis. (p<0.05 for all metabolites listed). *Indicates an increase with ETOM treatment.

**Table 2.**
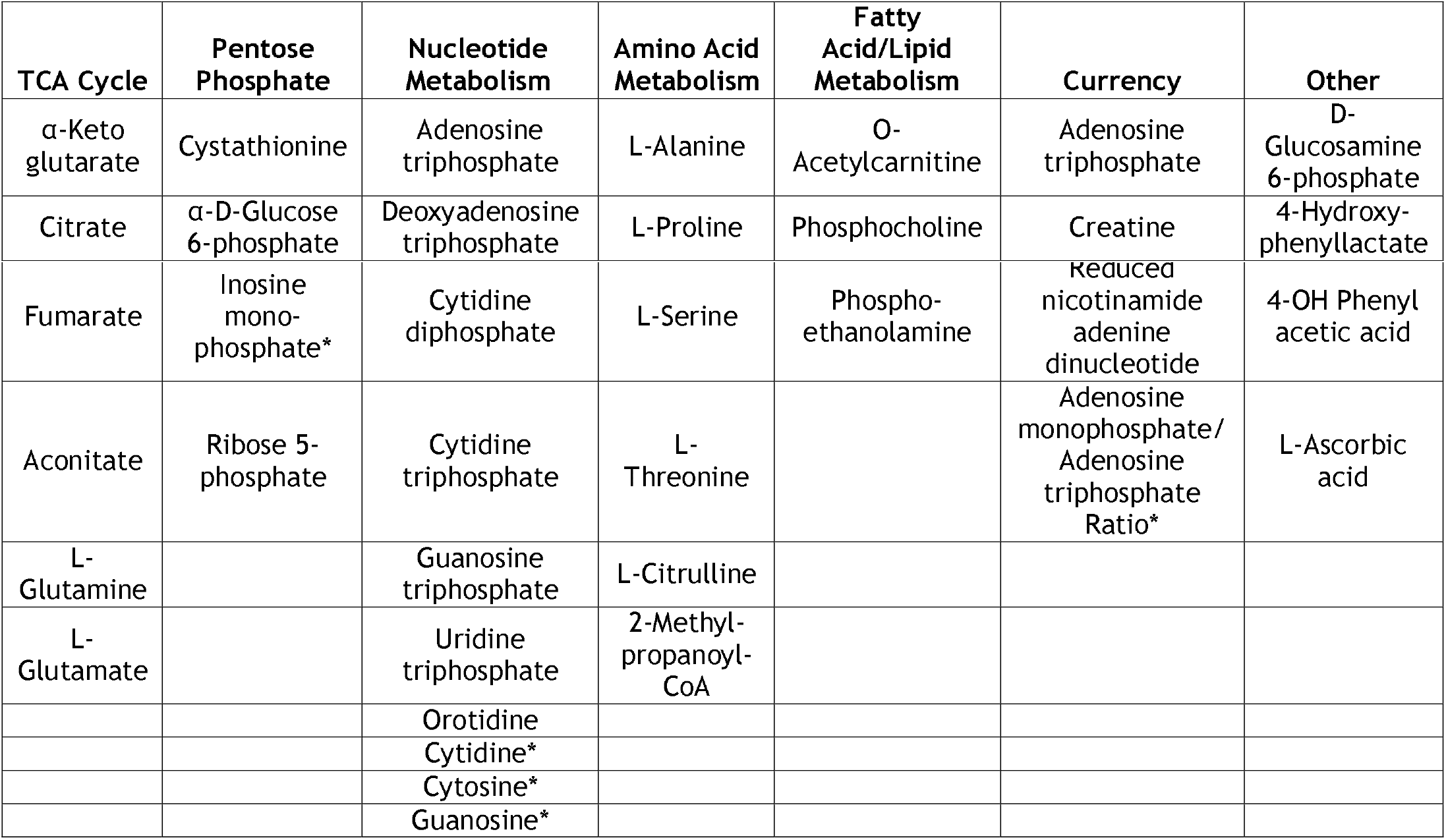
List of all metabolites altered by ETOM (100 μM) treatment with regard to relative amounts for HK157. Metabolites were measured by labeled ^13^C-palmitic acid using LC-MS analysis. (p<0.05 for all metabolites listed). *Indicates an increase with ETOM treatment.

### Fatty acid oxidation in IDH1 mutant GBM

Recent studies, including one from our own group (Garrett et al., 2018), have demonstrated that GBMs harboring an IDH1 mutation represent a metabolically distinct subset of glioma. Because of this, we sought to identify potential differences in FAO utilization between our IDH1 mutant and IDH WT gliomasphere cultures. Analysis of microarray data from 67 gliomasphere lines, 7 of which are IDH1 mutant, revealed elevated (though not statistically significant) mRNA expression of *CPT1A* in IDH1 mutant lines compared to IDH wildtype cultures (Figure 6A). Western blot analysis also showed significantly elevated CPT1A protein expression in three IDH1 mutant lines relative to wild type (Figure 6B). Interestingly, while there was a significant decrease in *CPT1C* mRNA expression in IDH1 mutants, we found no difference in protein expression (Figure 6A,B). Similar to IDH WT cells, LC-MS analysis with fully-labeled palmitic acid in an IDH1 mutant cell line, HK252, showed diminished labeling of acetyl-CoA and several TCA-associated metabolites when treated with ETOM (Figure 6C). Interestingly, 2-hydroxyglutarate (2-HG), a proposed oncometabolite (Dang et al., 2009) and the resulting product of the mutant IDH1 enzyme itself, was among the most heavily labeled metabolite in this patient line. Treatment with ETOM significantly diminished the fractional contribution suggesting that carbons from FAO are incorporated into the TCA, thus contributing to 2-HG production in IDH1 mutant GBMs. The relative amount of intracellular 2-HG is also reduced with ETOM. (Figure 6C-D). Interestingly, FAO inhibition with ETOM in IDH1 mutant cells did not result in a decrease in the relative amounts of TCA cycle intermediates as was observed in IDH WT cells (Figure S5), perhaps indicating a greater reliance on FAO to feed the TCA cycle in IDH WT GBM. Analysis of the full list of significantly different metabolites revealed fewer overall metabolites altered by ETOM treatment in IDH1 mutant cells (Table 3 shows fractional contribution and Table 4 shows relative amounts). These data confirm FAO utilization in both IDH1 mutant and wildtype cells while highlighting several potentially important metabolic differences between the two. Further investigation with a greater number of IDH mutant and wildtype lines will be needed to fully elucidate these differences.

**Figure 6.**
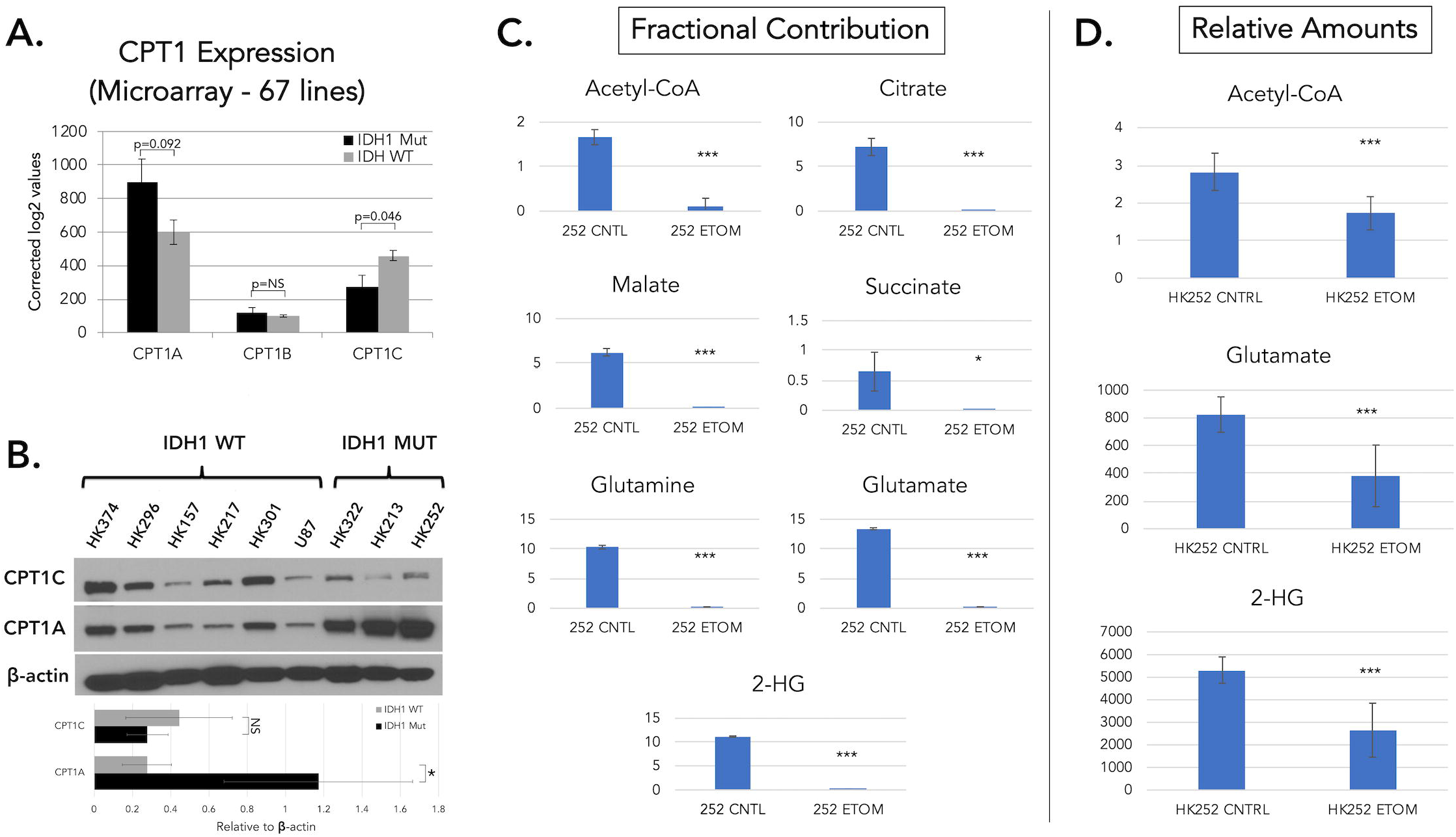
IDH1 mutant GBM cells express higher levels of CPT1A and utilize fatty acids to produce 2-HG. **A.** mRNA expression of *CPT1* isoforms in IDH1 mutant (N = 5) and wildtype (N = 62) patient-derived GBM cultures based on analysis of microarray data. **B.** Wwestern blot of CPT1A and CPT1C across a panel of IDH wildtype and IDH1 mutant lines. Error bars = ± SD. (* = p<0.05, Kruskal-Wallis). **C-D.** Analysis of the effects of ETOM (100 μM) in an IDH1 mutant cell line on fractional contribution and relative amounts of relevant metabolites using ^13^C-palmitic acid LC-MS. n = 3 replicates Error bars = ± SD. (* = p<0.05, ** = p<0.01, *** = p<0.001). See also Figure S5.

**Table 3.** List of all metabolites altered by ETOM (100 μM) treatment with regard to fractional contribution for IDH1 mutant HK252. Metabolites were measured by labeled ^13^C-palmitic acid using LC-MS analysis. (p<0.05 for all metabolites listed). *Indicates an increase with ETOM treatment.

| TCA Cycle | Nucleotide Metabolism | Amino Acid Metabolism | Fatty Acid/Lipid Metabolism | Currency | Other |
| --- | --- | --- | --- | --- | --- |
| Aconitate | Adenosine | L-Asparagine | O-Acetylcarnitine | Acetyl-CoA | N-Acetyl-D-glucosamine 6-phosphate |
| Citrate | Adenine | L-Aspartic acid | Cytidine diphosphate | Adenosine triphosphate |  |
| Malate | Adenosine triphosphate | L-Arginine* | Cytidine diphosphate ethanolamine | Reduced Glutathione |  |
| Succinate | Cytidine | Homocysteine |  | Oxidized Glutathione |  |
| 2-Hydroxyglutarate | Cytidine monophosphate | L-Proline |  |  |  |
| L-Glutamine | Cytidine triphosphate |  |  |  |  |
| L-Glutamate | Cytosine |  |  |  |  |
|  | Deoxyguanosine |  |  |  |  |
|  | Inosine |  |  |  |  |
|  | Orotidine |  |  |  |  |
|  | Uridine diphosphate |  |  |  |  |
|  | Uridine diphosphate glucose |  |  |  |  |
|  | UDP-N-acetylglucosamine |  |  |  |  |
|  | Uridine monophosphate |  |  |  |  |
|  | Uridine triphosphate |  |  |  |  |

**Table 4.** List of all metabolites altered by ETOM (100 μM) treatment with regard to relative amounts for IDH1 mutant HK252. Metabolites were measured by labeled ^13^C-palmitic acid using LC-MS analysis. (p<0.05 for all metabolites listed). *Indicates an increase with ETOM treatment.

| TCA Cycle | Nucleotide Metabolism | Amino Acid Metabolism | Currency | Other |
| --- | --- | --- | --- | --- |
| 2-Hydroxyglutarate | Adenine* | 2-Methylpropanoyl-CoA | Acetyl-CoA | D-Fructose 1,6-bisphosphate |
| L-Glutamate | Deoxyadenine* | Glycine | Creatine | α-D-Glucose 6-phosphate |
|  | Guanosine monophosphate |  | Reduced nicotinamide adenine dinucleotide | D-Glucosamine 6-phosphate |
|  | Inosine* |  |  | Pantothenate |
|  | Orotate* |  |  | Sorbitol |

## Discussion

Identifying metabolic vulnerabilities in cancer for use in targeted therapies has been a topic of interest for decades. While glucose metabolism has drawn the most attention in the scientific community, the ability of brain tumor cells to utilize other substrates for cellular maintenance and bioenergetics has become increasingly evident (Marie and Shinjo, 2011; Salzillo et al., 2016; Seyfried et al., 2011; Woolf et al., 2016). Several lines of evidence exist for the use of a ketogenic diet (KD) to treat brain tumors. The first lies with the innate ability of the human body to efficiently adapt to ketosis (Weber et al., 2018). For example, the KD has been successfully implemented to treat certain diseases such as refractory pediatric epilepsy with no clear sustained adverse health effects (Simeone et al., 2017; Youngson et al., 2017; Zhang et al., 2018). Additionally, several previous studies have indicated that many brain tumors lack the ability to oxidize ketones due to decreased expression of several mitochondrial genes (Seyfried et al., 2003; Skinner et al., 2009), although there is also evidence suggesting gliomas oxidize ketone bodies at comparable levels to the brain when exposed to a KD (De Feyter et al., 2016). Our study suggests that when glucose levels are diminished, GBM cells may adapt by partially shifting their metabolism toward ketone body and fatty acid oxidation. Prior animal studies investigating the KD have yielded mixed results, usually depending on the manner of administration and ability to achieve ketosis (Lussier et al., 2016; Meidenbauer et al., 2014; Stafford et al., 2010; Zhou et al., 2007). One study using U87 cells in a mouse tumor model found that an *ad libitum* KD increased blood ketone levels without affecting blood glucose or tumor growth (Zhou et al., 2007). Only with a calorie-restricted KD were glucose levels and tumor growth both diminished. This is consistent with studies demonstrating anti-tumor effects of calorie restriction irrespective of the KD (Jiang and Wang, 2013; Mukherjee et al., 2002; Seyfried et al., 2011). Here we report that an *ad libitum* KD significantly elevated blood ketone levels and decreased blood glucose, but resulted in a trend for increased tumor growth in the U87 line that was not statistically significant, and no effect on tumor growth in a patient-derived GBM line or in a syngeneic mouse tumor. Furthermore we found for all *in vivo* models used in this study the KD had at best no effect, or in the case of U87 cells, a significantly detrimental effect on mouse survival. While it is possible that the KD in GBM patients may have therapeutic utility, our data suggest that clinical trials investigating such treatments should consider the likelihood that brain tumors will be able to oxidize fatty acids for energy in the absence of additional treatments to modify this capability (Nebeling et al., 1995; Rieger et al., 2014; Schwartz et al., 2015; Zuccoli et al., 2010).

While also investigating ketone body utilization, this study primarily focused on the ability of GBMs to oxidize FAs. Building upon previous reports (Cirillo et al., 2014; Lin et al., 2017; Pike et al., 2011), our results demonstrate that both GBM cell lines and patient-derived GBM cell cultures express the necessary enzymes required for FAO, utilize FAs to support overall metabolism, and inhibition of this process has anti-proliferative effects. Importantly, our results also suggest FAO may be intrinsic to GBM and is not fully counteracted by AKT-enhancing mutations. Activation of the PI3K/AKT/mTOR pathway has been implicated in glucose addiction, a diminished ability to utilize alternative substrates that leaves malignant cells particularly vulnerable to glucose starvation (Elstrom et al., 2004; Maroon et al., 2015), and a metabolic shift towards FA synthesis (Guo et al., 2009). However, our current study suggests that under physiologically relevant glucose conditions (2.5 mM), FAs play a central role in the overall GBM metabolome regardless of mutational status. A study published during revision of the current manuscript found a difference in use of FAO in tumors based on TGCA classification (Kant et al., 2020). Specifically, FAO was found to be enhanced in mesenchymal tumor lines. However, we observed no such pattern with regard to TCGA subtype in our study. Our findings of a mutation-independent use of FAO may be due to methodological differences with previous studies. Furthermore, our *in vivo* findings that show significantly decreased survival and a trend towards increased U87 tumor growth with a KD support our *in vitro* findings that AKT-enhancing mutations do not make GBMs “glucose addicted”, as U87 cells themselves are PTEN deficient. The KD limits glucose availability to the tumor, so the enhanced tumor growth we observe is likely to have been supported by the use of alternative energy substrates despite U87s having PTEN mutations.

The utilization of FAO by these tumors may seem counterintuitive since the brain receives an ample supply of glucose, the most highly utilized substrate in GBM. ATP generation by FAO also demands more oxygen, is a slower overall process, and generates more superoxide than glycolysis (Schönfeld and Reiser, 2013), though it does produce more ATP than glycolysis. While the idea of simultaneous FA synthesis and oxidation seems counterproductive, studies such as ours make it increasingly clear that FAO makes malignant cells more resilient and adaptable to metabolic stress. FAs represent a readily available substrate that can be utilized to support both bioenergetic demands and for the production of other cellular components. Lipid levels in malignant gliomas are reportedly higher than in normal brain tissues (Srivastava et al., 2010). Gliomas can both produce free FAs and lipid stores endogenously via *de novo* synthesis from excess glucose and via uptake of exogenously produced free FAs from blood. Our experiments suggest that both endogenous and exogenous sources of FAs can be used as an energy substrate to support GBM cell growth. Perhaps the most notable aspect of FAO is that it results in the production of acetyl-CoA, an important metabolite that regulates a number of cellular processes including the TCA cycle. Recent evidence demonstrates that much of the acetyl-CoA produced by malignant gliomas is derived from sources other than glucose (Mashimo et al., 2014), highlighting FAO as a potentially critical component of brain tumor metabolism. In addition to ATP, FAO contributes to NADH and NADPH levels. While labeled carbon from FAO is found throughout the metabolome, it should not contribute net carbons to most metabolites, at least when even-chained FAs are oxidized, as for every two carbons of acetyl-CoA that are metabolized, two carbons are lost as carbon diaoxide. However, we found that inhibition of FAO resulted in the loss of relative amounts of a number of metabolites, in addition to fractional contributions, for reasons that are not yet clear, but which may be of use clinically. Interestingly, we found that inhibition of FAO resulted in the loss of relative amounts of some nucleotides, a finding previously reported for endothelial cells (Schoors et al., 2015).

A recent study reported that GBMs are reliant on FAO to sustain proliferation and that acyl-CoA binding protein (ACBP) promotes this process by regulating the availability of long chain fatty-acyl CoAs to the mitochondria (Duman et al., 2019). Our results provide further mechanistic insight by identifying CPT1A as a valid therapeutic target and demonstrating that inhibition of CPT1-mediated FAO decreased tumor growth and improved survival using a murine orthotopic xenograft model. In support of this idea, a recent study showed that systemic administration of ETOM in tumor bearing mice resulted in a trend toward decreased tumor growth and a significant increase in survival, both of which were amplified when combined with a gycolysis inhibitor (Kant et al., 2020). While the KD initiates production of ketone bodies, it also results in increased FA availability with the potential to be taken up and utilized by the tumor (Fraser et al., 2003; Taha et al., 2005). Because CPT1A knockdown decreased tumor growth from the KD, our results implicate FAs, rather than ketone bodies, as the primary metabolic substrates responsible for the growth-promoting effects of the KD. These results provide strong evidence that FAO is a significant pathway in GBM and may serve as a valuable target either alone or in combination with other currently investigated treatments.

Our study focused on the CPT1A isoform, and did not directly study the roles of the CPT1B and C isoforms. This was because the effects of etomoxir were largely mimicked by CPT1A knockdown. CPT1C is found in the brain and prior studies indicate that it plays important functional roles in tumors outside the brain (Chen et al., 2017; Sanchez-Macedo et al., 2013; Wang et al., 2018; Zaugg et al., 2011), although its role in gliomas is unknown and its precise functional role in FAO is also unclear. *CPT1B* is expressed at low levels in brain tumors, but it too could play a role in GBM FAO. A combinatorial knockout approach, while challenging, would shed light on the contributions of these other enzymes.

While investigating FA metabolism in GBM as a whole, we also sought to characterize potential differences between IDH1 mutant and wildtype GBM cells. Recent evidence suggests that GBMs harboring IDH mutations represent a distinct subclass of glioma (Cancer Genome Atlas Research Network et al., 2015). Our group previously demonstrated IDH1 mutant gliomaspheres are metabolically distinguishable from their WT counterparts based on expression profiles, glucose consumption, and nucleotide synthesis utilization (Garrett et al., 2018). The results of the current study suggest further metabolic differences with regard to FAO. Analysis of CPT1 isoform expression revealed a trend for increased expression of CPT1A in IDH1 mutant cultures relative to IDH wildtype that was not statistically significant. LC-MS tracing experiments with ^13^C-palmitic acid revealed extensive FA utilization throughout the metabolomes of both IDH1 mutant and WT cells. However, the effects of FAO inhibition with ETOM were more pronounced within the TCA cycle of IDH WT cells. Importantly, 2-HG was among the most heavily labeled metabolites in IDH1 mutant cells. Treatment with ETOM significantly decreased the amount of 2-HG, indicating that biproducts of FAO may serve as a vital component for the production of this oncometabolite. While these effects were only tested in a single IDH1 mutant cell line, and thus require further investigation, our results suggest FAO represents an additional mechanism contributing to the unique metabolic phenotype of IDH mutant gliomas.

Although our findings emphasize the complex and highly adaptable nature of brain tumor metabolism, important aspects of these studies require further investigation. The preponderance of evidence reported in this study support the conclusion that GBMs oxidize FAs. Inhibition of FAO resulted in overall deficits in cellular growth and FA-mediated mitochondrial respiration. However, while FA supplementation under physiologically relevant glucose concentrations demonstrated inverse effects to that of inhibition in some cell lines, including U87, this effect was not consistent across all patient-derived cell lines tested, suggesting some variability in the ability of GBMs to utilize exogenous FA. Our LC-MS studies also clearly show that FAs are taken up, oxidized, and utilized throughout the GBM metabolome. Finally, many of the effects of FAO inhibition achieved with ETOM were validated using multiple methods of CPT1A knockdown, indicating that they are not due to off-target effects of ETOM. The fact that ETOM has been shown to exhibit off-target effects in certain cells at high doses cannot be overlooked, and could ultimately limit its use clinically. Developing more potent and specific FAO inhibitors, could prove essential for targeting brain tumors in patients. However, due to the grim prognosis associated with glioma and the currently inadequate treatment options available, this study highlights several important aspects of brain tumor metabolism that have direct application to improving current standard of care therapy.

### Limitations of the Study

The provenance of the U87 cell line as been lost, and therefore may not be representative of patient GBMs, although it has been verified as a glioblastoma cell line (Allen et al., 2016). This limitation is somewhat mitigated by our observations in a large array of primary GBM lines *in vitro* and one primary GBM line *in vivo.* A modestly high concentration of ETOM (100μM) was used for experiments described, allowing the possibility for off-target effects of the drug, which were mitigated at least partially by the observation that the effects of ETOM were largely mimicked by CPT1A knockdown. Our data suggest that CPT1A is the isoform primarly responsible for FAO in GBM, however we do not know the extent of potential compensation from the B and C isoforms when CPT1A is knocked down. The ketogenic diet used in our *in vivo* models was calorie-unrestricted. While we did observe a decrease in blood glucose and an increase in ketones, there might be more of an effect on tumor growth in an unrestricted ketogenic diet model. Additionally the timeframe for the ketogenic diet in these experiments was limited due to the fast rate of growth of the tumors. A longer-term ketogenic diet, such as that which a tumor patient would undergo, might be more effective at slowly tumor growth as cellular building blocks become more scarce over time. The metabolimics of FAO inhibition are described in three cell lines (U87, HK157, HK252) and therefore may not be representative of all GBM.

## Resource Availability

Lead Contact: Further information and requests for resources and reagents should be directed to and will be fulfilled by the Lead Contact, Harley I. Kornblum, M.D. Ph.D.

## Materials Availability

No new or unique materials were generated by this study.

## Data and Code Availability

The raw data for the U87 metabolomics presented in Figure 2 can be found at DryadData.org titled “U87_Isotopologues_ETOM_CPT1A_CRISPR”, https://doi.org/10.5068/D1R95W.

## Acknowledgements

This work was supported in whole or part by National Institute of Health (NIH) Grants: NS052563 (H.I.K.) and CA179071 (A.L.) the UCLA SPORE in Brain Cancer (P50 CA211015); and the Dr. Miriam and Sheldon G. Adelson Medical Research Foundation (H.I.K.). Thanks go to the UCLA Institute for Digitical Research and Education for statistical consultation.

## Author Contributions

Jantzen Sperry: Conceptualization, Methodology, Formal Analysis, Investigation, Writing - Original Draft, Writing - Reviewing & Editing,

Michael C. Condro: Conceptualization, Methodology, Formal Analysis, Investigation, Writing

Reviewing & Editing,

Lea Guo: Investigation, Methodology,

Daniel Braas: Methodology, Investigation, Data Curation, Writing - Original Draft, Writing - Reviewing and Editing,

Nathan Vanderveer-Harris: Investigation,

Kristen K.O. Kim: Investigation,

Whitney B. Pope: Supervision, Writing - Reviewing & Editing, Funding Acquisition,

Ajit S. Divakaruni: Conceptualization, Methodology, Writing - Reviewing & Editing, Validation,

Albert Lai: Conceptualization, Writing - Reviewing & Editing,

Heather Christofk: Conceptualization, Supervision, Writing - Reviewing & Editing,

Maria G. Castro: Resources, Writing - Reviewing & Editing,

Pedro R. Lowenstein: Resources, Writing - Reviewing & Editing,

Janel E. Le Belle: Conceptualization, Methodology, Validation, Writing - Original Draft, Writing - Reviewing & Editing, Supervision,

Harley I. Kornblum: Conceptualization, Resources, Writing - Original Draft, Writing - Review & Editing, Supervision, Project Administration, Funding Acquisition.

