## Supplemental Files for "Glioblastoma utilizes fatty acids and ketone bodies for growth allowing progression during ketogenic diet therapy"

**Supplementary Materials:**

**Supplemental Figures**

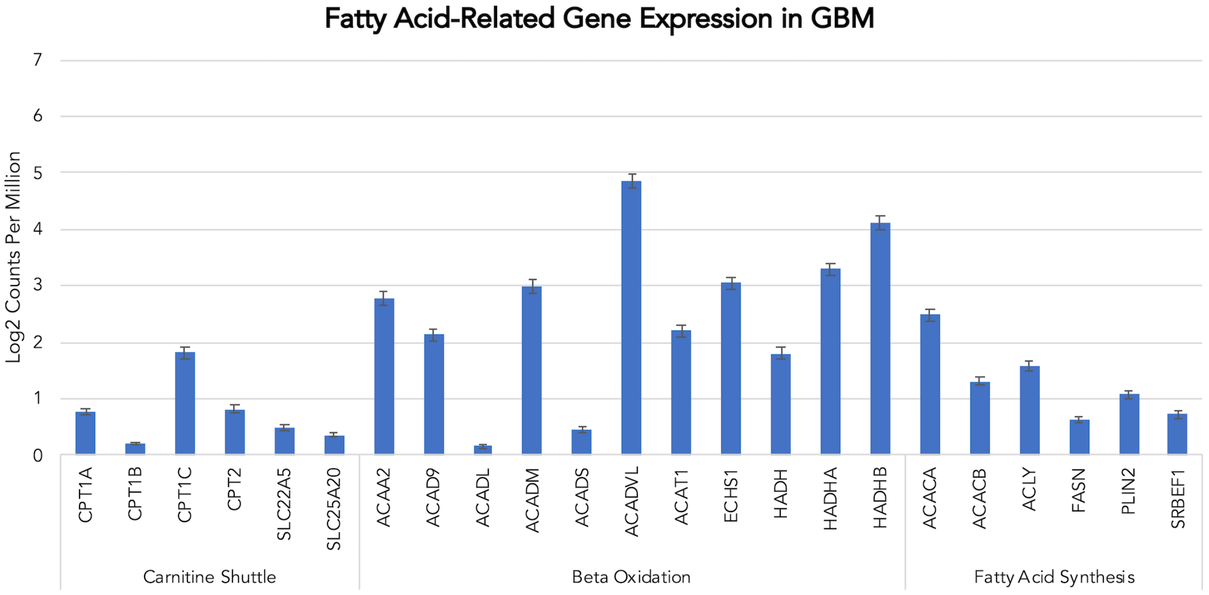


**Figure S1: Expression of Fatty Acid-Related Genes in GBM, Related to Figure 1A.**

Average expression of genes related to mitochondrial FAO and synthesis by single cell RNAseq from four patients. Data represent expression levels from neoplastic cells found either at the tumor core (n=1029 cells) or in the periphery (n=62 cells). Error bars ± SEM.

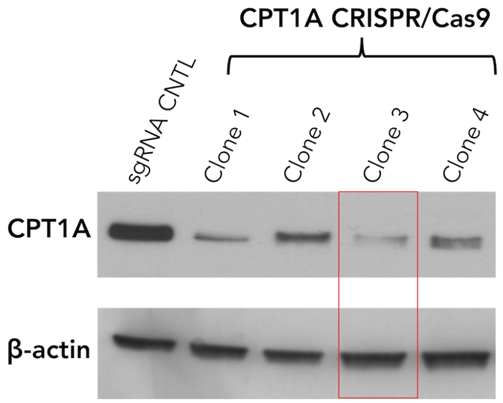


**Figure S2: Efficiency of CRISPR/Cas9 targeting CPT1A, Related to Figure 2.**

Western blot of lysates from U87 cells expressing Cas9 and either control sgRNA or one of 4 CPT1A-targeting sgRNA clones. Due to its increased efficiency at reducing CPT1A, clone 3 was used in the experiments shown in Figure 2.

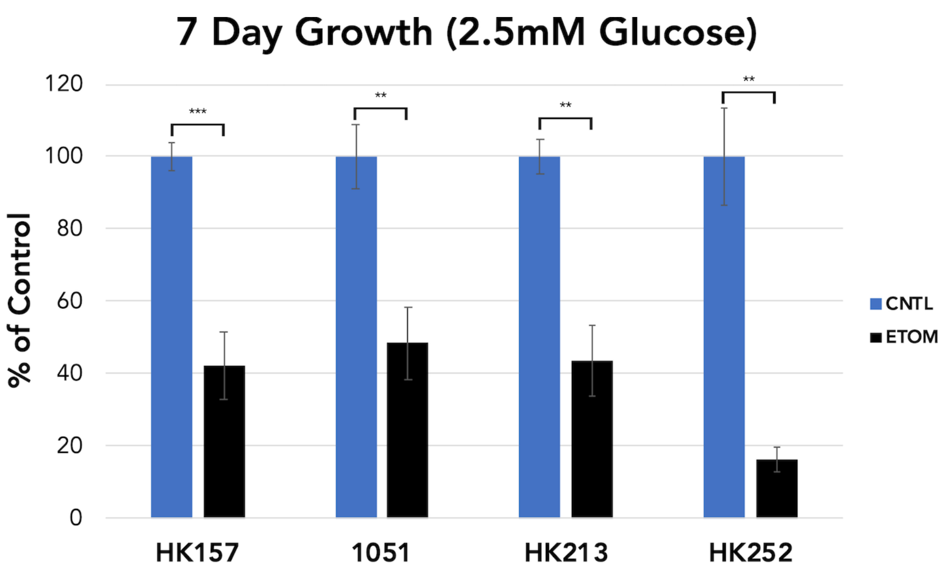


**Figure S3: ETOM inhibits growth of patient-derived GBM cell lines in 2.5 mM glucose, but response to FA or 3-OHB supplementation varies, Related to Figure 4A.**

7 day growth assessment (cell counts) with 100 µM ETOM, 50 mM palmitate (FA), and 1.25 mM 3-OHB in 2.5 mM glucose. n = 3 replicates. Error bars = ± SD. (* = $p<0.05$, ** = $p<0.01$, *** = $p<0.001$).

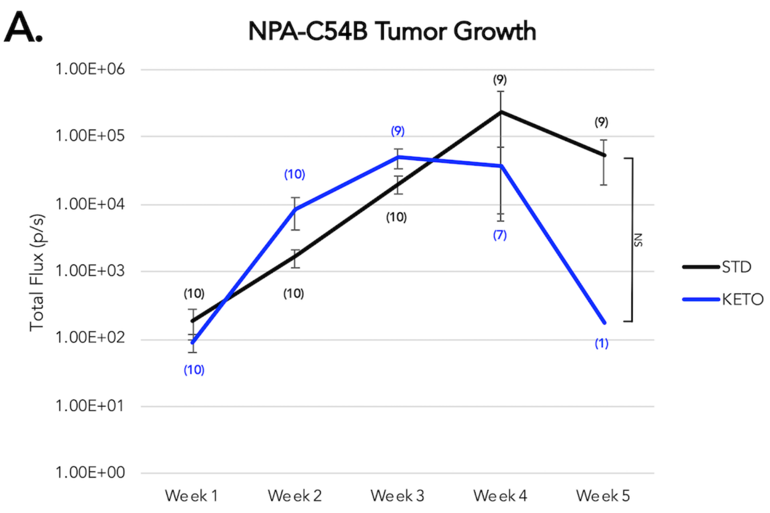


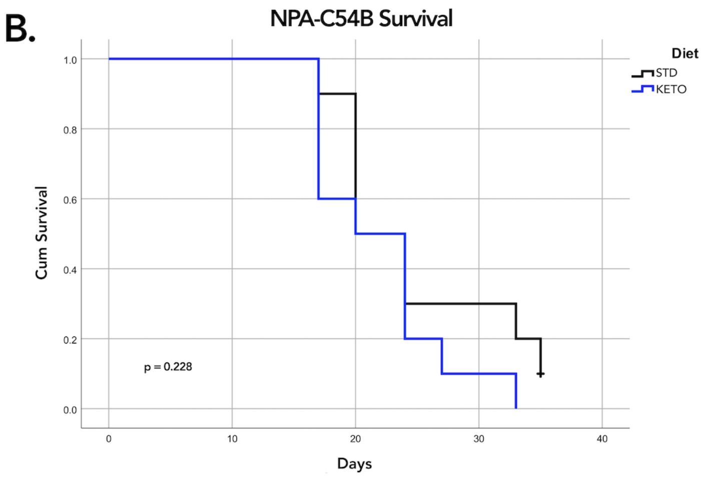


**Figure S4: The unrestricted ketogenic diet yields no benefit to tumor growth or survival in a mouse model of GBM, Related to Figure 4E and 4F.**

Tumor growth (left) measured by luciferase imaging and survival (right) of a murine model of GBM (NPA-C54B) implanted into immune-competent C57Bl/6 mice. Numbers in parentheses indicate the number of mice averaged for each timepoint. Error bars = ± SEM. (* = p<0.05, ** = p<0.01, *** = p<0.001).

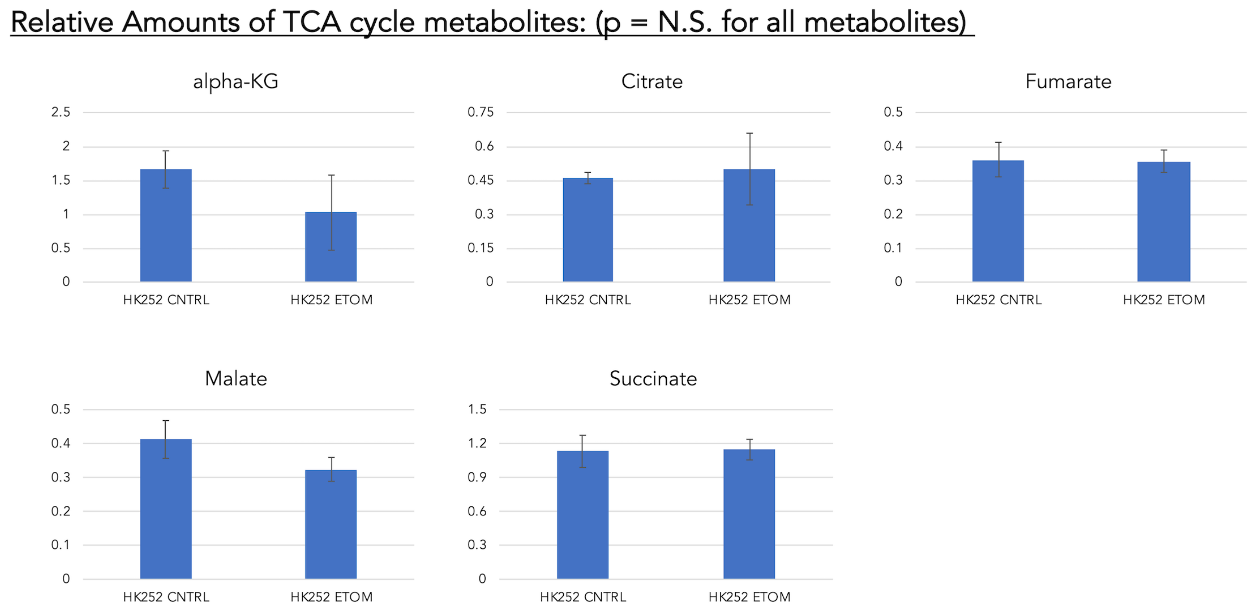


**Figure S5: CPT1 inhibition with ETOM does not alter relative amounts of TCA metabolites in IDH1 mutant cells, Related to Figure 6.**

Analysis of the effects of 100 µM ETOM on relative amounts of TCA metabolites in an IDH1 mutant line using $^{13}C$-palmitic acid (LC-MS). n = 3 replicates. Error bars = ± SD. p>0.05 (N.S.) for all metabolites shown.

### Transparent Methods:

#### Collection and Maintenance of *In Vitro* Cultures

Tumor samples were collected under institutional review board-approved protocols and graded by neuropathologists. Patient-derived gliomasphere cultures were prepared as follows: immediately after receiving resected tissue, samples were digested with papain and acellular debris was removed. The remaining cells were incubated in gliomasphere defined media containing DMEM/F-12 supplemented with B27, heparin, EGF, bFGF, and penicillin/streptomycin until sphere formation was achieved. Gliomasphere stocks were frozen down at approximately passage 5 to maintain cells at low passage throughout the study. All cell lines used in this study, including U87, were continuously grown in serum-free gliomasphere media.

#### Expression Analysis

Relevant expression, mutational status (such as EGFR and IDH1), and patient characteristics for the majority of cell lines used in this study have been previously reported (Laks et al., 2016; Garrett et al., 2018). The following cultures are known to be PTEN deficient: U87, HK217, HK229, HK296, HK301, but there may be other cultures that are PTEN mutated/deleted as well. Also of note, while BT142 is IDH1 mutant (R132H), it does not produce 2-HG (Luchman et al., 2013). HK211 is both IDH1 (R132H) and EGFRviii mutant. Known status of EGFR, PTEN, and IDH1 are indicated in Figure 6A for patient-derived GBM cultures. Analysis of *CPT1* isoform expression is based on the microarray dataset (GSE98995) previously described by Laks et al. 2016 and available in the Gene Expression Omnibus (GEO) repository.

#### Variable Glucose Media and Substrate Supplementation

All media described contain B27, heparin, EGF, bFGF, and penicillin/streptomycin. For 25 mM glucose the base media used is Neurobasal-A Medium ([-] Glutamine) (Gibco #10888-022) supplemented with GlutaMax Supplement (Gibco #35050061). 2.5 mM and 1 mM glucose media was made by combining the Neurobasal-A Medium ([-] Glutamine) above with Neurobasal-A Medium ([-] D-glucose, [-] Sodium Pyruvate) (Gibco #A24775-01) to achieve the desired glucose concentration. Glutamax Supplement and sodium pyruvate (Gibco #11360-070) were then added to reach a final concentration of 1X. Palmitic acid was purchased from Sigma-Aldrich (#P0500) and conjugated with fatty acid free, low endotoxin bovine serum albumin (BSA, Sigma Aldrich, #A1595). Carnitine (Sigma Aldrich, #C0283) was included in fatty acid-supplemented conditions. 3-hydroxybutyrate (3-OHB) was also purchased from Sigma (#166898).

**shRNA and CRISPR/Cas9 Lentiviral Knockdown**

Plasmids with shCPT1A (RHS4531-EG1374 glycerol set) and the non-silencing shCNTL (RHS4346) cloned into pGIPZ expression vectors were purchased from Dharmacon. The firefly luciferase-GFP virus (FLuc-GFP; backbone= pRRL-sinCMV-iresGFP) was produced by UCLA Vectorcore and supported by Molecular Technologies Core (IMTC) CURE/P30 DK41301-26. These plasmids were transfected into 293T cells along with 2$^{nd}$ generation viral ΔR8.74 package and VSV-g envelope for production of lentivirus.

**U87 In Vivo Orthotopic Xenotransplants**

All animal experimentation was performed with institutional approval following NIH guidelines using adult NSG mice. Intracranial xenotransplants were performed similarly to previous descriptions with minor changes (Laks et al., 2009).

*CPT1A /Ketogenic Diet:*

$10^5$ U87 cells expressing either shCNTL or shCPT1A along with firefly luciferase-GFP (FLuc-GFP) were stereotactically injected unilaterally into the striatum of NOD *scid* gamma (NSG) mice under isoflurane anesthesia over 5 minutes using the following coordinates: 1.5 mm lateral and 0.5 mm anterior to bregma, and 3.0 mm below the pial surface. N=10 mice per group. As shown in Figure 3B, mice were kept on standard diet for 4 days to allow appropriate recovery, after which half of each group was placed on the ketogenic diet (BioServ, S3666). On Day 8, weekly bioluminescence imaging was initiated as described below. On Day 21 mouse weight, blood glucose and ketone measurements were taken. Tumor growth was measured until each animal either died or became moribund. Mouse survival was tracked for all groups.

*HK408 and NPA-C54B :*

Transplants and institution of the ketogenic diet for this experiment were carried out as described above with the exception that there was no knockdown of CPT1A group and therefore only 20 total mice were used (10 per group). For HK408, $10^5$ cells were implanted into 10 male and 10 female NSG mice, and groups were divided evenly by sex. No sex differences were found with regards to the ketogenic diet (data not shown). In the murine tumor model, $5x10^3$ cells of NPA-C54B were implanted into the striatum of 20 male C57Bl/6 mice as described in Nuñez et al., 2019, at coordinates 1.0 mm anterior and 2.5 mm lateral to bregma, and 2.5 mm deep. These cell lines were chosen because of their consistency in forming tumors that grow quickly. The speed of tumor formation was important, not simply for convenience, but it allowed us to take into account concerns regarding significant weight loss and the overall health of animals receiving a long-term ketogenic diet. As above, mice were kept on a standard diet for four days after surgery, at which time they were divided into groups of standard and ketogenic diet. Tumor growth was measured by weekly bioluminescence imaging as described above, and survival was tracked until mice died or became moribund.

**Bioluminescent Imaging**

Optical imaging was performed at the Preclinical Imaging Technology Center at the Crump Institute for Molecular Imaging at UCLA. 100 µl of D-luciferin (GoldBio) dissolved in phosphate buffer saline without Ca2+ or Mg2+ (30 mg/ml) was introduced to each animal by intraperitoneal (IP) injection. After 7 minutes of conscious uptake, mice were anesthetized by inhalation of 2.6% isoflurane in oxygen and placed in dedicated imaging chambers. The IVIS Lumina 2 imaging system (Caliper Life Sciences) was utilized for *in vivo* bioluminescent imaging. Luminescence was measured over 3 minutes, and a corresponding photograph of the mice was taken and co-registered with the luminescent image for signal localization. Images were analyzed by drawing regions of interest and quantified as total flux (photons/second) with the Living Image software package (Perkin Elmer).

**Western Blot and Immunohistochemistry**

Western blots were performed using the following antibodies: mouse anti CPT1A (abcam, 128568), rabbit anti CPT1C (LsBio, LS-C167010), rabbit and mouse anti beta-actin (Abcam). To prepare samples for immunohistochemical analysis, Mouse brains from U87 transplant experiments were perfused using 4% paraformaldehyde (PFA) and incubated overnight in PFA at 4°C for 24 hours. Tissue was washed with PBS and incubated in 20% sucrose at 4°C for a minimum of 24 hours in preparation for sectioning on a cryostat. Sections (20 µm thick) were post-fixed for 15 minutes with cold 4% paraformaldehyde followed by 3 washes with TBS prior to performing immunohistochemistry for CPT1A.

**Acute etomoxir treatment and fatty acid and ketone supplementation**

Dissociated cultures were plated at a density of $5x10^4$ cells/ml in triplicate in gliomasphere defined media containing 25 mM glucose unless otherwise indicated. Cells were incubated for 24 hours after which point 100 µM etomoxir (ETOM) ((+)-Etomoxir sodium salt hydrate, Sigma Aldrich, #E1905) was added to each sample and again on day 4. For FA and ketone supplementation, $4x10^3$ cells were plated per well in 96-well plates and incubated for 24 hours, after which either FAs or ketones were added to yield the following final concentrations of each. For FA: 25-200 µM palmitate (Sigma Aldrich, #P0500) bound to fatty-acid free BSA (5g/50mL in 10% PBS) (Sigma Aldrich, #A1595) and 500 µM carnitine (Sigma Aldrich, # C0283). For 3-OHB (Sigma Aldrich, #166898) final concentrations were 1.25-10 mM. Blood plasma concentrations of these substrates in healthy adults range from 111-260 µM palmitate and 67 µM carnitine (Borch et al., 2012; Cunnane et al., 2012; Jensen et al., 1989). Cells were allowed to grow for the indicated period of time prior to analysis using the Dojindo Cell Counting Kit 8 (#CK04-20) according to manufacturer instructions.

**Cell growth analysis**

Single-cell suspensions were made from bulk cultures of the U87 and patient-derived glioma cell lines and counted using a Countess automated cell counter. Cells were plated at $5x10^4$ cells/mL in 6-well plates and grown as gliomaspheres in control or experimental conditions. For

ETOM experiments, cells were treated 24 hours after initial plating and treatment was re-administered on Day 4. After 7 days the gliomaspheres in each well were fully dissociated and re-counted to assess changes in cell number during treatment. This analysis of cell number examines both cell survival and proliferation in response to treatment.

**Proliferation analysis with bromodeoxyuridine (BrdU)**

Single cell suspensions were plated as a monolayer on laminin-coated glass coverslips (Sigma, L2020) in 24-well plates followed by a 4 day ETOM treatment. Two BrdU pulses were given 2 hours apart and cells were fixed for 15 minutes with 4% paraformaldehyde. Coverslips were washed with PBS and immunocytochemistry was performed to assess the percentage of BrdU-positive cells by visual microscopic cell counts.

**Annexin V/PI Flow Cytometry Analysis**

Single cell samples were treated with 100 μM ETOM for 4 days after which spheres were re-dissociated with 200 μl accumax, centrifuged, and washed with PBS. Annexin V/propidium iodide (PI) staining was carried out according to manufacturer protocol using the Annexin V APC flow cytometry kit (Thermo Fisher) while including a 1 μl/ml final concentration of PI. Samples were gently mixed and incubated with Annexin V/PI binding buffer and incubated for 15 minutes at room temperature protected from light. Samples were kept on ice and analyzed within 1 hour by flow cytometry.

**LC-MS with fully labeled $^{13}C$ palmitate**

Gliomaspheres were dissociated into single cells with Accumax™ and $2x10^5$ cells were cultured for 48 hours in 2.5 mM glucose media in triplicate for each sample. Cells were then rinsed with PBS and re-plated in 2.5 mM glucose media, either unlabeled or containing 200 μM fully-labeled $^{13}C$-palmitic acid for an additional 48 hours. Cells were then centrifuged and rinsed with 1ml ice-cold 150 mM ammonium acetate (pH 7.3). Centrifugation was performed again and 1ml of ice-cold 80% methanol was added. Cells were transferred to an Eppendorf tube, and 10 nmol norvaline (Sigma-Aldrich, N7502) was added to each sample. Samples were centrifuged for 5 min at top speed and the supernatant was transferred into a glass vial. Samples were resuspended in 200 μL cold 80% methanol, followed again by centrifugation, after which the supernatant was added to the glass vial. Samples were dried in an EZ-2Elite evaporator. The remaining pellet was resuspended in RIPA buffer and a Bradford assay was performed to quantify total protein concentration for sample normalization. Dried metabolites were resuspended in 50% ACN and 5 μL loaded onto a Luna 3 μm NH2 100 A (150 × 2.0 mm) column (Phenomenex). The chromatographic separation was performed on an UltiMate 3000 RSLC (Thermo Scientific) with mobile phases A (5 mM NH4AcO pH 9.9) and B (ACN) and a flow rate of 200 μL/min. The gradient from 15% A to 95% A over 18 min was followed by 9 min isocratic flow at 95% A and re-equilibration. Metabolite detection was achieved with a Thermo Scientific Q Exactive mass spectrometer run in polarity switching mode (+3.5 kV/− 3.5 kV). TraceFinder 4.1 (Thermo Scientific) was used to quantify the area under the curve for metabolites by using accurate mass measurements (< 3 ppm) and the retention time of purchased reference standards. Relative amounts of metabolites were calculated by summing up all isotopologues of a given metabolite and normalized to cell number. Correction for naturally occurring $^{13}C$ as well as calculation of fractional contributions and clustering analyses were done in R. Fractional contribution was calculated as $\sum_{i=1}^{n} \frac{M_i \cdot i}{n}$, where n is the number of carbons in the metabolite, $i$ is the iteration of each possible $^{13}C$-labeled carbon, and $M_i$ is the relative abundance of the $i^{th}$ isotopologue. The relative amount is calculated as the sum of all isotopologues of each metabolite normalized to total protein.

**Statistical Analysis**

Statistical analysis was performed using either Microsoft Excel, GraphPad Prism, or IBM SPSS software with guidance from the UCLA Institute for Digital Research & Education. For tumor growth, flux data was lognormalized and statistical significance was determined using a mixed effects model with repeated measures for individual animals and Tukey's post-hoc pairwise analyses. Survival was analyzed by Kaplan-Meier curves with pairwise Mantel-Cox post-hoc analyses. For other comparative samples, normality of distributions was determined by Shapiro tests and significance was determined using either ANOVA or a univariate generalized linear model model and two-tailed Student's T-tests where appropriate. All quantitative data and associated error bars represent the mean ± either the standard deviation (SD) or standard error of the mean (SEM) as indicated for each figure. Experiments were performed in triplicate at a minimum, with 95% confidence intervals and p values calculated for relevant comparison. For all figures, p values are represented as follows: NS = not signifant, * = $p<0.05$, ** = $p<0.01$, *** = $p<0.001$.
